# Cost-effective epigenetic age profiling in shallow methylation sequencing data

**DOI:** 10.1101/2021.10.25.465778

**Authors:** Alexandre Trapp, Vadim N. Gladyshev

## Abstract

There is a critical need for robust, high-throughput assays of biological aging trajectories. Among various approaches, epigenetic aging clocks emerged as reliable molecular trackers of the aging process. However, current methods for epigenetic age profiling are inherently costly and lack throughput. Here, we leverage the *scAge* framework for accurate prediction of biological age from very few bisulfite sequencing reads in bulk samples, thereby enabling drastic (100-1,000-fold) reduction in sequencing costs per sample. Our approach permits age assessment based on distinct assortments of CpG sites in different samples, without the need for targeted site enrichment or specialized reagents. We demonstrate the efficacy of this method to quantify the age of mouse blood samples across independent cohorts, identify the effect of calorie restriction as an attenuator of the aging process, and discern rejuvenation upon cellular reprogramming. We propose that this framework may be used for epigenetic age prediction in extremely high-throughput applications, enabling robust, large-scale and inexpensive interventions testing and age profiling.

## Introduction

Mammalian aging is characterized by profound changes in the epigenomic landscape, particularly in the DNA methylome^1, 2^. Cytosine methylation fractions at CpG dinucleotides can be detected using next-generation sequencing approaches—both genome-wide and targeted—as well as microarray technologies^3–7^. These methods permit high-resolution characterization of methylation patterns in bulk tissue samples, and more recently in single cells^8, 9^. While methylation plays an active role in a variety of downstream biological processes, its relationship with aging has been of special interest in recent years. In fact, methylation dynamics have been integrated with artificial intelligence techniques (including both shallow and deep machine learning approaches) to enable accurate age estimation based on DNA methylation profiles in humans, mice, and other mammalian organisms^7, 10–15^.

Epigenetic aging clocks, tools that quantitatively leverage coordinated age-associated changes in the DNA methylome, are believed to reflect an inherent measure of biological age in their predictions^16^. While chronological age is fixed based upon one’s date of birth, biological age is conceived of as an integrative and fluid metric that accounts for intrinsic and environmental factors, which are ultimately reflected in the dynamic DNA methylome. Epigenetic age, specifically measured by tracking concerted changes in particular CpG methylation patterns, has hence emerged as one of the most promising “biomarkers” of the aging process, and has been used as a platform to evaluate the effect of longevity and rejuvenation interventions^17, 18^. This line of inquiry is especially promising, given that epigenetic aging dynamics have recently been shown to exhibit remarkable conservation across mammalian species^19–22^.

However, current methods for epigenetic age profiling are much too costly and low-throughput. In humans, the most common approach for partial methylome analysis remains the various suite of methylation arrays produced by Illumina (including the depreciated 27K & 450K chips, and the more recent EPIC Infinium chip)^7, 23^. These microarrays leverage bisulfite conversion and fluorescent probes that ultimately enable reproducible assessment of the fraction of methylation in a bulk sample (known as the beta values) at several thousand CpGs^24^. In mice, the lack of methylation arrays (until recently^19, 25^) has restricted profiling to reduced representation bisulfite sequencing (RRBS), a method that involves restriction-enzyme-mediated CpG island enrichment, bisulfite conversion, and short-read next-generation sequencing^11, 18^. With both of these approaches, there is a requirement for considerable amounts of input DNA, in part due to harsh chemical treatment by sodium bisulfite during library preparation. Additionally, the costs (either of the microarray or the sequencing runs) can be prohibitively high for routine applications. This poses a profound limitation for large-scale efforts to profile biological age in populations or cohorts, particularly in regard to throughput, labor, time and cost.

To address these issues, we leveraged our recently developed *scAge* framework^26^ to enable robust profiling of epigenetic age in low-pass bulk bisulfite sequencing data. By downsampling deeply-sequenced RRBS data, we arrived at a data type very similar to single cells: sparse, mostly binary methylation values. The use of few sequencing reads further results in variable coverage of cytosines, despite the use of enriched methylation protocols. To address this, our method makes use of different CpG sites in each sample for epigenetic age predictions, circumventing the need for a set of CpGs to be consistently covered across many samples. Here, we introduce our framework as a tool for inexpensive and high-throughput epigenetic age predictions, particularly well-suited for large-scale studies.

## Results

### Simulating low-pass bisulfite sequencing data

To develop a platform for epigenetic age profiling from low-coverage bisulfite sequencing data, we first designed a simulation pipeline that enables the creation of reduced methylation profiles from randomly downsampled sequencing reads (Fig. 1a). We utilized existing RRBS data from mouse aging studies^10, 27^, wherein individual libraries were deeply sequenced at more than 10^7^ reads per sample. Next, we devised a random downsampling algorithm that takes as input individual reads and outputs a desired number of subsampled reads. Practically, we downsampled bulk data to 10^2^, 10^3^, 10^4^ and 10^5^ reads, enabling comprehensive assessment of different low-pass bisulfite sequencing outputs.

**Figure 1:**
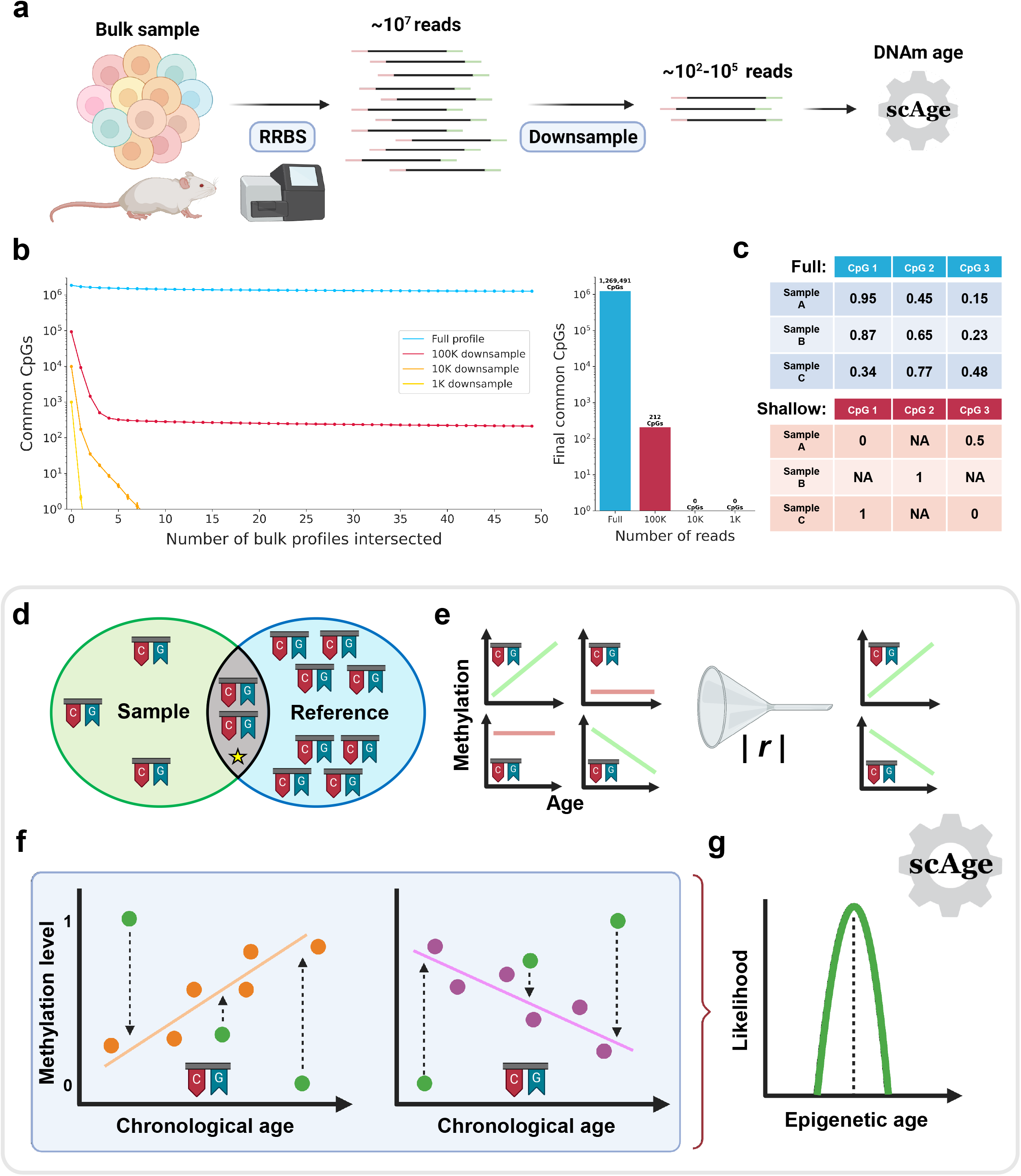
An approach for low-pass epigenetic age profiling. a) Schematic of the analysis pipeline. Murine bulk samples were collected and RRBS libraries were prepared and sequenced to a high depth (>10^7^ reads). Processed methylation data was downsampled with a reproducible random seed to 10^2^-10^5^ reads, resulting in limited CpG profiles for each subsample. Lastly, epigenetic age (“DNAm age”) was predicted by a modified version of the *scAge* framework. b) Scatterplot (left) and barplot (right) representing the number of common CpGs after progressive intersections of full RRBS profiles as well as downsampled profiles (100K, 10K, and 1K reads) in the Thompson *et al.*^27^ blood data. In the scatterplot, individual dots show the median of 100 permutations for each intersection, with each permutation producing a randomized intersection order. Y-axis is log-scale. Full RRBS profiles produce many intersections (blue), while downsampled profiles produce minimal to no intersections. Error bars depicting 95% confidence intervals are shown, but in most cases are smaller than the size of the points. The barplot shows the final number of common CpGs for a particular profile size. c) Schematic tables highlighting the results of intersecting full RRBS profiles (blue, top) or downsampled profiles (red, bottom). With full profiles, many CpGs are covered consistently and at high depth across samples, resulting in valid fractional values within the unit region. However, with shallow (low-pass) sequencing, fewer reads are assessed in each sample, resulting in discordant CpG coverage, sparsity, and a primarily binary data type. d) Intersection step of low-pass *scAge*, wherein only CpG sites common to both a sample and the reference dataset (highlighted by the star in the center) are retained for downstream processing. e) Filtration step of low-pass *scAge*, wherein only common CpG sites exhibiting a robust positive/negative linear relationship between methylation level and age are chosen for downstream processing. f) Probability computation step of low-pass *scAge*. Deeply sequenced samples are used as training data to construct linear regression models mapping changes in methylation as a function of chronological age. Individual orange/purple dots schematically depict training samples, and orange/purple lines depict ordinary least squares regressions based on these samples. Probability computations were performed by subtracting from 1 the distance from a particular observed CpG site methylation level in shallow data (green) to the linear regression estimate. g) Likelihood distribution generated for a sample based on the collective probabilities harnessed across many CpGs. The dashed line depicts the age of maximum likelihood, which is interpreted as the epigenetic age of a particular sample.

Since bulk RRBS protocols commonly enable readout at a few million CpGs per sample, downsampling to a low number of reads or performing low-pass sequencing limits the applicability of conventional epigenetic clock approaches. These methods traditionally rely on training machine learning models—usually based on elastic-net regularization—which effectively select informative CpGs and assign them weights in a resulting linear model. Subsequently, new methylation values, weighted by the model and often adjusted by an intercept, can be directly used for epigenetic age predictions^7, 10, 12, 27–29^. However, the algorithms driving these models require that many CpGs are covered at high depth consistently across many samples, which can presently only be accomplished by deep sequencing or methylation array technologies. Additionally, once a set of CpG sites is chosen and given weights in the model, the same CpG sites must all be present in any future testing dataset in order to obtain the most accurate predictions. This can be circumvented to some degree with imputation approaches^12, 27, 30, 31^, but an excessive number of missing values tends to greatly reduce the absolute accuracy of current clocks^32^. While intersecting bulk genomic RRBS data produces large feature tables (on the order of 10^5^-10^6^ CpGs) amenable to machine learning approaches, low-pass sequencing profiles have minimal common sites, precluding the use of conventional methods (Fig. 1b). Indeed, feature tables constructed from aggregated shallow data are a stark contrast to those built from deep data, specifically differing in their primary data type and their sparsity (Fig. 1c).

To address this key challenge, we leveraged the *scAge* framework, which is particularly amenable to sparsely covered methylation data with distinct sets of CpG sites available in each individual sample. Briefly, the framework first involves training CpG-specific regression models predicting methylation level from chronological age. Next, our platform employs a ranked intersection algorithm: only CpG sites covered both in a particular sample of interest and the reference data are retained, and these are subsequently ranked and filtered to output only highly age-associated common CpG sites (Fig. 1d, e). The algorithm then cycles through each CpG site in the resulting profile, and measures the distance between the observed methylation value and the estimate from the linear model at a particular age. We propose to interpret this distance metric as a probability, meaning that ages with smaller absolute distance between the observed value and the model prediction are more representative of the true biological age of that particular sample (Fig. 1f). Finally, we amass several hundred/thousand CpG-specific probabilities into a comprehensive likelihood profile, which enables efficient epigenetic age profiling using sparse methylation data (Fig. 1g).

The *scAge* framework was initially developed to profile epigenetic age using high-resolution single-cell methylation data^26^. Here, we adjusted the original algorithm to conform to an additional parameter resulting from its application to bulk samples: meaningful non-binary methylation. In single cells, the binary nature of DNA methylation data is mostly reflective of inherent biology in major cell types. However, bulk samples (even downsampled ones) may exhibit meaningful non-binary methylation, given that sequencing reads come from many different cells in the sample. In line with this notion, we modified our algorithm to accommodate for non-binary methylation data. Altogether, we developed a pipeline that can take as input real or simulated low-pass methylation data for efficient epigenetic age predictions.

As reference data for *scAge*, we used normal, standard *ad libitum* C57BL/6J mouse blood samples from the Petkovich *et al.*^10^ (n = 153) and Thompson *et al.*^27^ (n = 50) studies. Samples ranged in age from 1 month to 35 months in the Petkovich *et al.* data, and 2 to 21 months in the Thompson *et al.* data. We selected highly covered (depth ≥ 5x) CpG sites in each sample and constructed a large feature table for all the samples in a dataset across all identified CpGs. Upon additional filtering, this generated two independent reference sets: 1.2 million CpG-specific linear models in the Thompson *et al.*^27^ data and 1.9 million CpG-specific linear models in the Petkovich *et al.*^10^ data. Of note, several differences in data processing, experimental protocols, and the number of samples in each dataset likely account for the variability in the number of trainable CpG linear models we obtained for each dataset.

### scAge tracks the aging process in blood based on low-pass sequencing data

We first trialed our approach on downsampled data from the Thompson *et al.*^27^ study (Fig. 2a). We selected 10^2^, 10^3^, 10^4^ and 10^5^ reads, each across 5 random states; in effect, this enabled randomization of the particular set of CpGs in each subsample, and mirrored sequencing small random aliquots of libraries. For each of these random, downsampled CpG methylation matrices, we applied our modified *scAge* framework trained on all samples from either the Petkovich *et al.*^10^ or Thompson *et al.*^27^ studies. We benchmarked the framework and its predictions across several read number parameters and likelihood profile sizes, using anywhere from 50 to 10^4^ top age-associated CpGs per sample for age computations.

**Figure 2:**
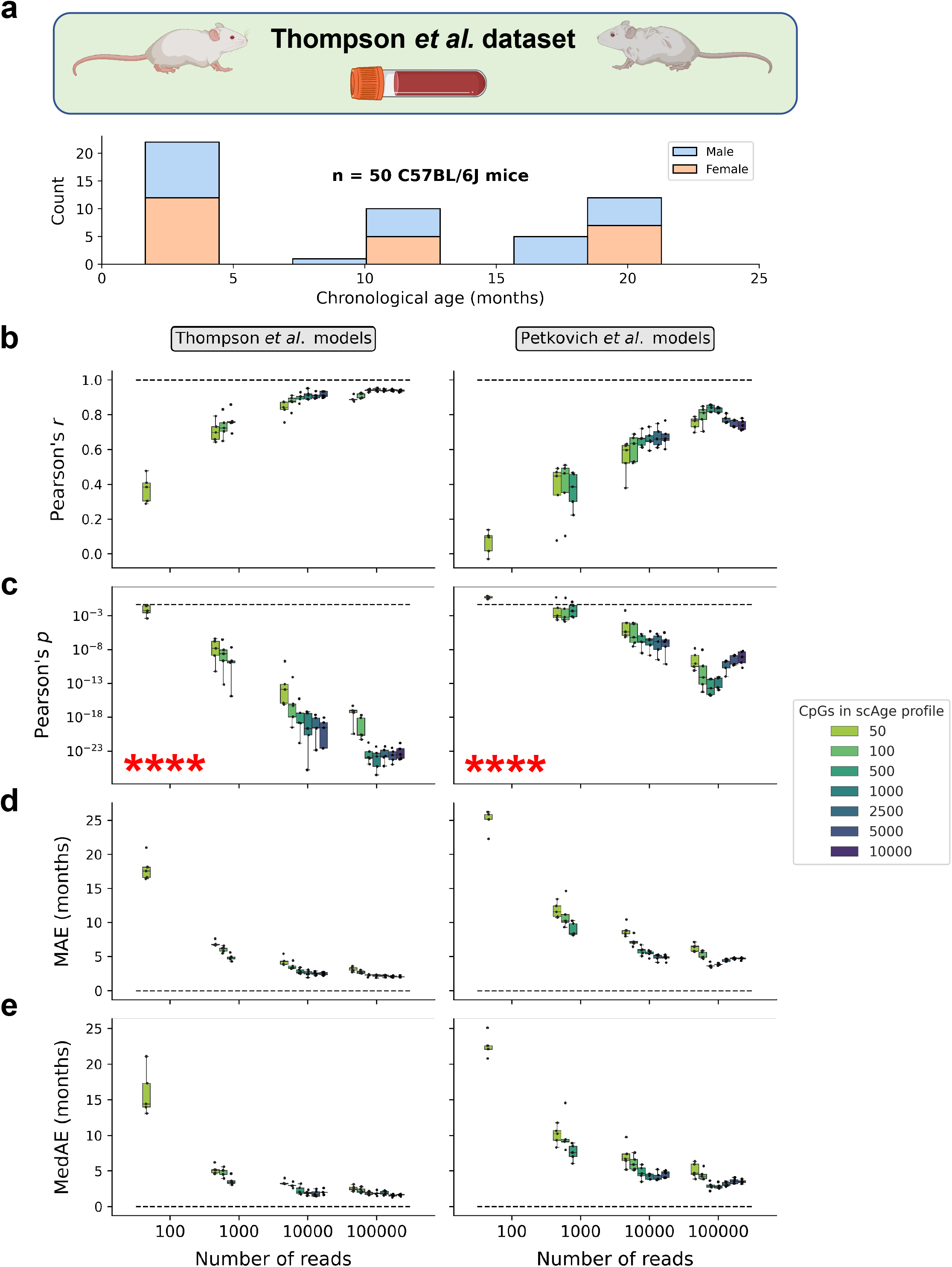
Low-pass epigenetic age profiling of blood data from Thompson *et al*. a) Schematic of the dataset (top) and age/sex distribution (bottom) of blood samples from C57BL/6J mice. Females (n = 24) are shown in orange, and males (n = 26) in blue. b-e) Boxplots of prediction metrics on the entirety of the Thompson *et al.* dataset. Predictions using Thompson *et al*. reference models are shown in the left panels, and predictions harnessing regression models built with the Petkovich *et al.* data are shown on the right. Individual dots (black) depict prediction metrics for a particular set of: 1) a random state 2) a number of downsampled reads, and 3) an *scAge* profiling parameter (the number of CpGs included in the age computation likelihood profile). Individual boxplots depict the median and the 1^st^ and 3^rd^ quartiles, with whiskers extending to 1.5x the interquartile range. Boxplots are colored based on the number of CpGs to profile in the likelihood computation. (b) depicts Pearson correlation coefficients, (c) depicts the associated two-tailed p-values, (d) depicts the mean absolute errors in months, and (e) depicts the median absolute errors in months. Red asterisks (****) highlight that significant p-values below 0.0001 were reached. Dashed lines highlight *r* = 1, p = 0.05 (nominal significance), MAE = 0, and MedAE = 0.

Excitingly, we observed robust performance of our epigenetic age profiling approach in these data, with the results varying based on the number of total CpGs in the downsampled data, the number of age-associated CpGs included in the likelihood profile, and the particular reference models employed. Pearson correlation coefficients (*r*) generally increased in magnitude and significance as more reads were subsampled, with some variation based on the size of the likelihood profile (Fig. 2b, c). Mean and median absolute errors decreased using both models as the number of reads was increased, again with some minor variation depending on profile sizes (Fig. 2d, e). Using these benchmarking results, we conducted further analysis on the best performing set of parameters according to both training datasets (100,000 reads with the top 1,000 age-associated CpGs included in the likelihood computation profile). When Thompson *et al.*^27^ linear models were used as reference, correlation coefficients ranged from 0.93-0.96 depending on the random seed (Extended Data Fig. 1). Application of the independent Petkovich *et al.*^10^ models showed slightly decreased predictive accuracy, with Pearson correlations ranging from 0.81-0.85. However, median errors were remarkably low with both training datasets, ranging from 1.81-2.35 months with Thompson *et al.* references models and 2.56-3.48 months with Petkovich *et al*. reference models. Interestingly, predictive metrics were almost equally robust when only 10,000 reads per sample were assayed (Fig. 2b-e). This suggests that integration of the *scAge* framework and low-pass sequencing—producing a relatively small number of reads—may be sufficient to accurately profile biological age.

Next, we applied the same methodology on downsampled data from the Petkovich *et al.*^10^ study, which included 153 blood samples from standard *ad libitum* male C57BL/6J mice aged 1-35 months (Fig. 3a). Again, we observed strong performance across both reference models on these data, with similar evaluation trends as what we observed upon application to downsampled Thompson *et al.*^27^ data (Fig. 3b-e). Indeed, increasing the number of reads was associated with improved prediction accuracy across all metrics, and results varied based on the specific number of CpGs leveraged by *scAge*. When selecting the most accurate parameters based on benchmarking (100,000 reads per sample with the top 500 age-associated CpGs included in the likelihood profile), Pearson *r* coefficients ranged from 0.88-0.9, and median absolute errors ranged from 2.9 to 3.2 months using the Petkovich *et al*. reference models. However, the Thompson *et al.* models were noticeably less accurate, particularly in regard to absolute error (*r* = 0.71-0.74, MedAE = 7.0-7.6m) (Extended Data Fig. 2). We hypothesize this may be largely due to differential processing of the Thompson *et al*.^27^ methylation data as compared to the Petkovich *et al.*^10^ data, and may also be reflective of batch effects. Indeed, we observed only a moderate positive association between CpG-specific Pearson correlations (*r* = 0.43) or linear regression coefficients (*r* = 0.54) with age among the Petkovich *et al.* and Thompson *et al.* reference datasets (Extended Data Fig. 3).

**Figure 3:**
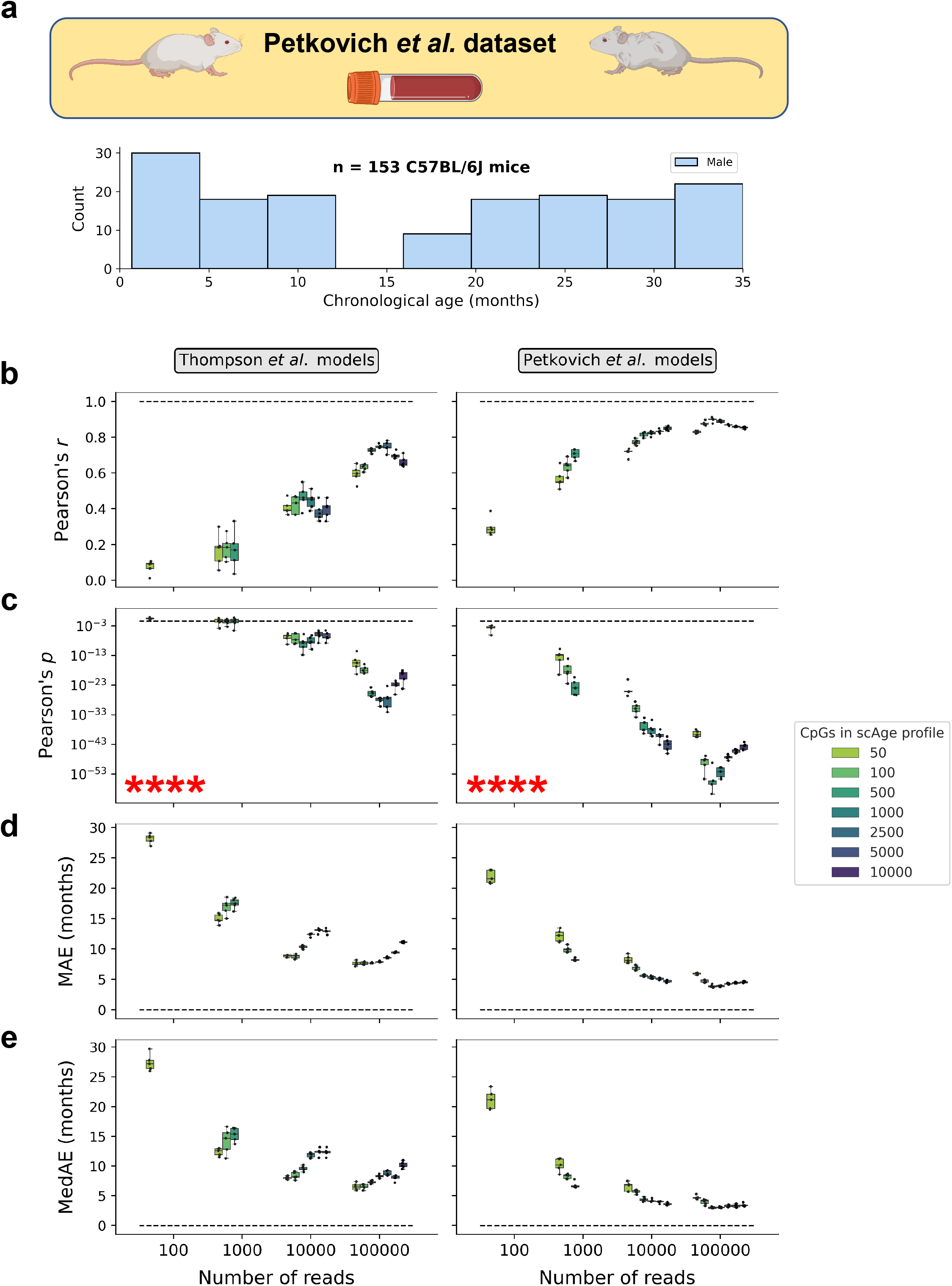
Low-pass epigenetic age profiling of blood data from Petkovich *et al*. a) Schematic of the dataset (top) and age distribution (bottom) of blood samples from male C57BL/6J mice (n = 153). b-e) Boxplots of predictive metrics on the entirety of the Petkovich *et al.* dataset. Predictions using Thompson *et al*. reference models are shown in the left panels, and predictions harnessing regression models built with the Petkovich *et al.* data are shown on the right. Individual dots (black) depict prediction metrics for a particular set of: 1) a random state 2) a number of downsampled reads, and 3) a *scAge* profiling parameter (the number of CpGs included in the likelihood profile). Individual boxplots depict the median and the 1^st^ and 3^rd^ quartiles, with whiskers extending to 1.5x the interquartile range. Boxplots are colored based on the number of CpGs to profile in the likelihood computation. (b) depicts Pearson correlation coefficients, (c) depicts the associated two-tailed p-values, (d) depicts the mean absolute errors in months, and (e) depicts the median absolute errors in months. Red asterisks (****) highlight that significant p-values below 0.0001 were reached. Dashed lines highlight *r* = 1, p = 0.05 (nominal significance), MAE = 0, and MedAE = 0.

### Prediction consistency suggests reproducibility in low-pass epigenetic age profiling

In order to effectively simulate different sequencing runs of the same sample, we applied a random downsampling approach that returned distinct sets of CpGs depending on the chosen seed. This procedure can be thought of as computationally analogous to picking a random aliquot of a library for sequencing, and is particularly suited for tracking the consistency of predictions from different subsamples (Fig. 4a). Excitingly, the best performing predictive tests (those with 10,000-100,000 reads) showed really strong inter-subsample prediction consistency. Indeed, both Thompson *et al.* and Petkovich *et al.* models had inter-subsample Pearson correlation coefficients greater than 0.9 when testing on downsampled Thompson *et al.*^27^ data (Fig. 4b). The Petkovich *et al.*^10^ models similarly showed robust inter-subsample prediction correlations when applied to Petkovich *et al.*^10^ data, while the Thompson *et al.*^27^ models applied to this same dataset had slightly weaker correlations (Fig. 4c). Overall, these results suggest that our approach is able to generate reproducible predictions from different small assortments of covered CpGs, reinforcing the putative value of our framework for handling shallow data.

**Figure 4:**
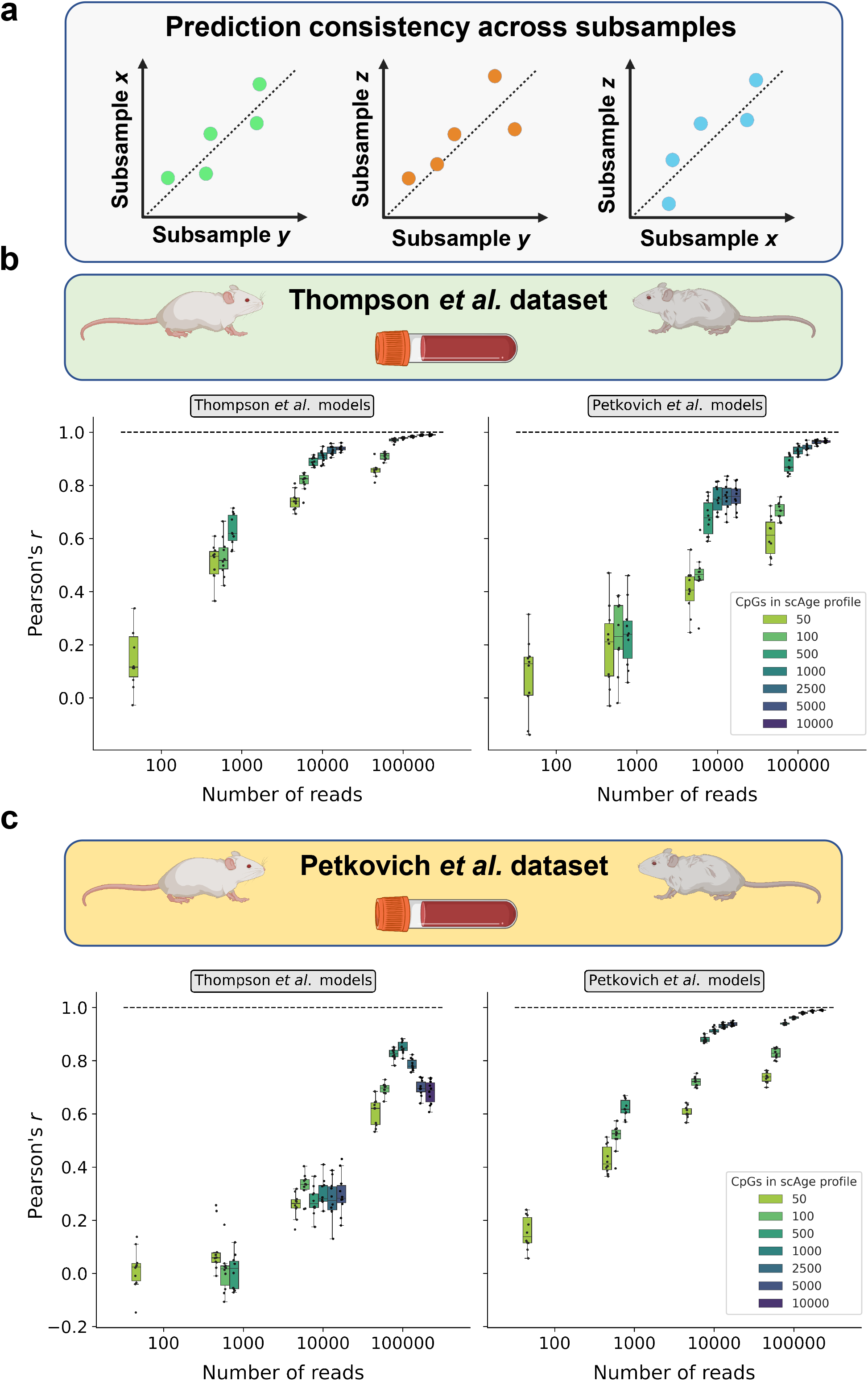
Consistency of low-pass epigenetic age predictions across random subsamples. a) Schematic of the analyses presented in this figure. Epigenetic age predictions from different random subsamples are compared, and the linearity of their relationship is evaluated by Pearson correlation analysis. b-c) Boxplots of inter-subsample Pearson correlation coefficients (*r)* on the entirety of (b) the Thompson *et al.* dataset and (c) the Petkovich *et al.* dataset. Inter-subsample associations using the Thompson *et al*. reference models are shown in the left panels, and inter-subsample associations using the Petkovich *et al.* models are shown on the right. Individual dots (black) depict Pearson correlations between predicted epigenetic age across two different random seeds for the same sample; since 5 random seeds were utilized in total, 10 possible combinations of random seeds are possible (n = 10 dots/boxplot). Individual boxplots depict the median and the 1^st^ and 3^rd^ quartiles, with whiskers extending to 1.5x the interquartile range. Boxplots are colored based on the *scAge* parameter used for these predictions (i.e., the number of CpGs to profile in the likelihood computation). The dashed lines in the boxplots highlight *r* = 1.

### Validation of attenuated aging with caloric restriction using low-pass epigenetic age profiling

Next, we were interested if our low-pass *scAge* framework could distinguish the effect of caloric restriction as a longevity intervention. Caloric restriction was previously found to attenuate biological aging as measured in blood^10^ and liver^13^ samples, and is commonly considered one of the gold-standard lifespan-extending interventions currently in use in model systems^33, 34^. To test the effectiveness of our approach, we analyzed 20 blood samples from male calorie-restricted C57BL/6J mice with chronological ages ranging from 10 to 27 months (Fig. 5a). Of note, these samples were completely independent from the Petkovich *et al.* linear model reference set used for *scAge* predictions.

**Figure 5:**
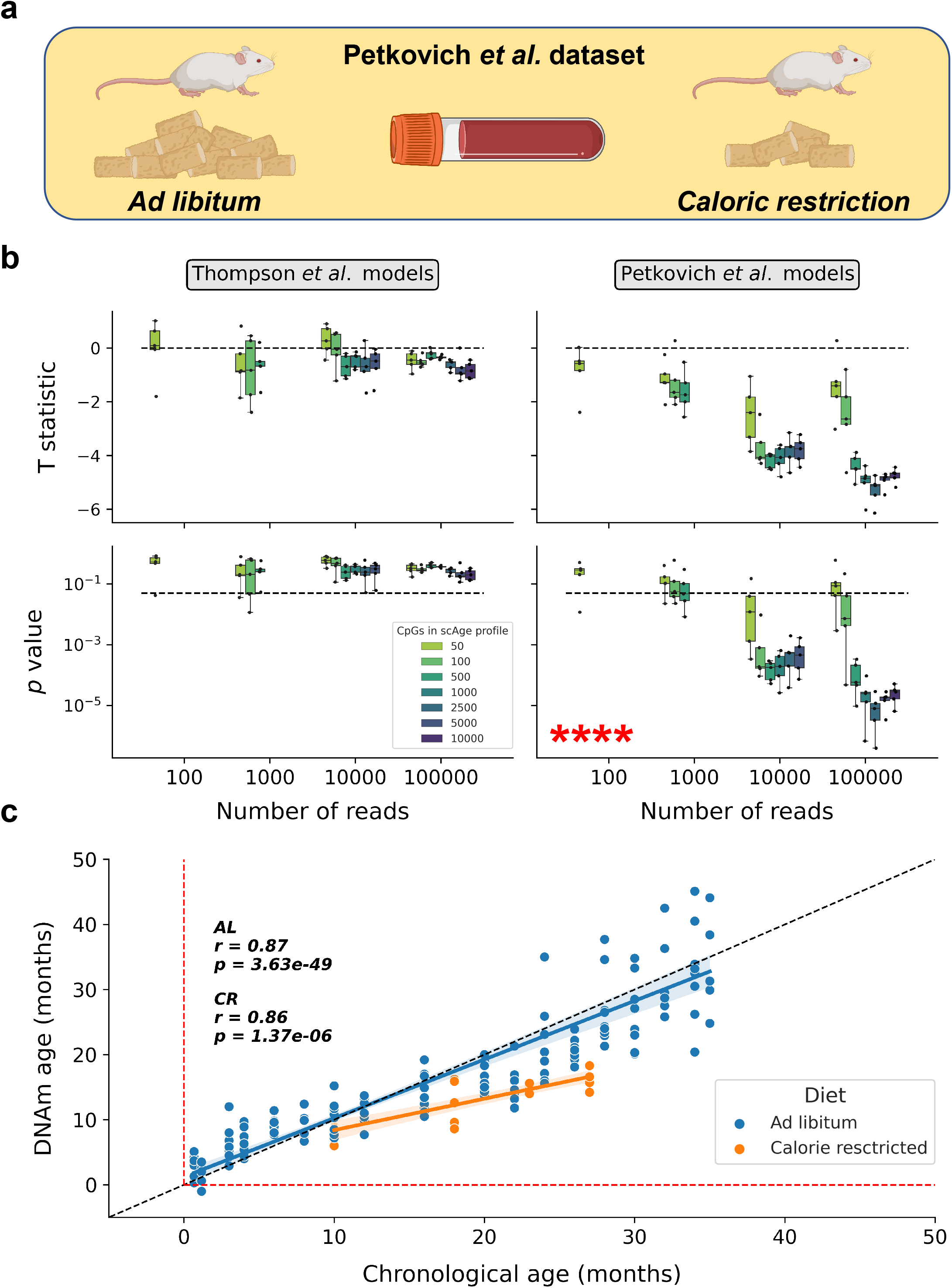
Attenuated epigenetic aging trajectories in response to calorie restriction. a) Schematic outline of the analyses presented in this figure. Blood age predictions and trajectories in standard *ad libitum* (n = 153) and calorie restricted (n = 20) male C57BL/6J mice originating from the Petkovich *et al*.^10^ study were compared. b) Boxplots of statistical testing metrics based on delta age measurements (epigenetic age – chronological age). Significance testing metrics using the Thompson *et al*. reference models are shown in the left panels, and those using the Petkovich *et al.* models on the right. Individual dots (black) depict prediction metrics for a particular random state, number of downsampled reads, and *scAge* profiling parameter. Boxplots show the median and the 1^st^ and 3^rd^ quartiles, with whiskers extending to 1.5x the interquartile range. Boxplots are colored based on the *scAge* parameter used for these predictions (i.e., the number of CpGs to profile in the likelihood computation). Upper panels depict the T statistic from Welch’s t-test used to quantify statistical significance between delta age in *ad libitum* and calorically restricted (CR) samples, with negative values indicating lower delta age in CR samples. Lower panels depict the one-tailed p-values associated with these t-tests. Red asterisks (****) highlight that significant p-values below 0.0001 were reached. The dashed lines in the upper panels highlight T = 0, and the dashed lines in the lower panels highlight nominal significance (p = 0.05). c) Scatterplot showing the relationship between epigenetic age and chronological age for *ad libitum* mice (blue, n = 153) and calorie-restricted mice (orange, n = 20) for one random state with 100,000 downsampled reads and the top 2,500 age-associated CpGs included in the likelihood profile. The dotted black line represents the identity line between chronological and DNAm age. Red dotted lines highlight DNAm and chronological ages of 0 months. The Pearson correlation coefficient (*r)* and its associated two-tailed p-value is shown separately for each group in the upper left corner. Regression models (blue/orange) are shown with 95% confidence intervals (light blue/orange).

Remarkably, by using these standard-diet Petkovich *et al.*^10^ models integrated into *scAge*, we were able to reliably detect a significant decrease in delta age (i.e., the difference between epigenetic and chronological age) for calorically restricted mice compared to control when either 10^4^ or 10^5^ reads per sample were utilized (Fig. 5b). In the best performing test case based on benchmarking (100,000 reads per sample with the top 2,500 CpGs included in the likelihood profile), delta age in calorically restricted mice was significantly (*p* < 0.0001) lower than in mice fed *ad libitum* across every random subsample (Extended Data Fig. 4). Moreover, linear regression analyses revealed a significantly attenuated epigenetic aging trajectory for calorie-restricted mice (Fig. 5c). This suggests that our method is capable of reproducibly discerning the effect of potent longevity interventions, despite having relatively few sequencing reads. However, it is important to note that application of the Thompson *et al*. models did not reveal a significant effect. Again, we hypothesize this is largely a manifestation of impaired accuracy of the Thompson *et al.* models on the Petkovich *et al.* data as a whole (Fig. 3b-e), which may be caused by processing incompatibilities, protocol differences, or other significant batch effects (Extended Data Fig. 3).

### Low-pass scAge identifies age reversal effect upon iPSC reprogramming

Given the rising interest in cellular reprogramming approaches for rejuvenation research, we were interested if our low-pass approach, combined with the *scAge* framework, could identify a significant epigenetic age decrease resulting from iPSC reprogramming^18, 35, 36^. We applied our method, trained on blood methylation data, to renal and lung fibroblasts and corresponding iPSC lines derived from these tissues. Despite a rather small sample size (n = 3 cell lines per group), we observed that predicted epigenetic ages based on Petkovich *et al.* blood regression models were significantly lower for iPSCs than for fibroblasts across all random subsampling states tested when using 100,000 reads (Fig. 6). When selecting the best test case based on benchmarking (100,000 reads per sample with the top 500 CpGs included in the likelihood profile), we witnessed significant decreases in epigenetic age across almost all random subsamples using both the Petkovich *et al.* and Thompson *et al.* reference models, in both kidney-derived and lung-derived iPSCs (Extended Data Fig. 5, 6). Absolute age predictions had improved accuracy under Petkovich *et al.* models, with iPSC samples displaying epigenetic ages near 0, validating results from earlier deep sequencing, elastic-net based approaches^7, 32^.

**Figure 6:**
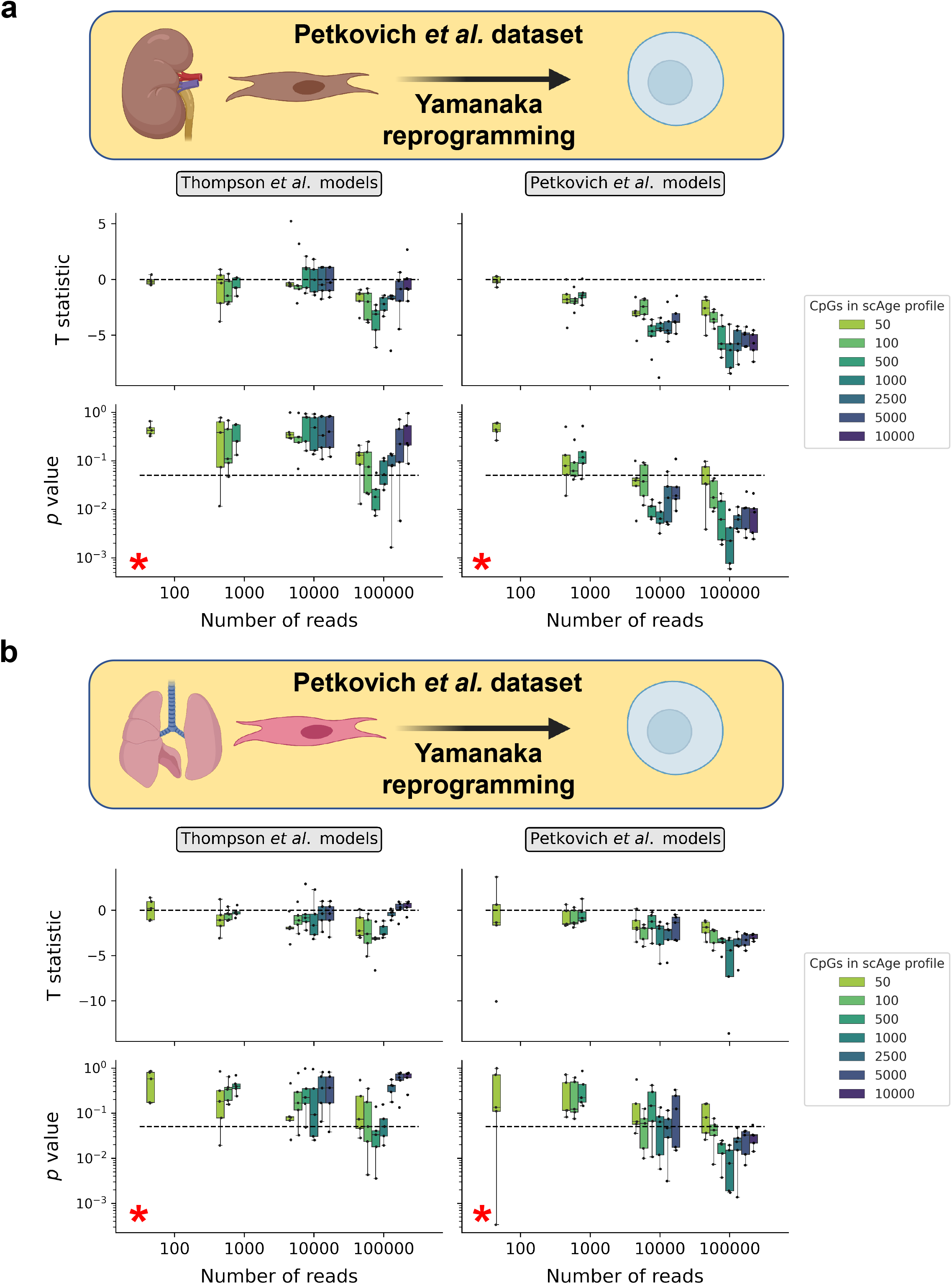
Low-pass *scAge* tracks epigenetic age reversal by iPSC reprogramming. a-b) Experimental schematics (top), and boxplots (bottom) of statistical testing metrics based on epigenetic age measurements in (a) renal fibroblasts and iPSC samples derived from these cells and (b) lung fibroblasts and iPSC samples derived from these cells, all from the Petkovich *et al.*^10^ study (n = 3 per group). Significance testing metrics using the Thompson *et al*. reference models are shown in the left panels, and those using Petkovich *et al.* reference models are shown on the right. Individual dots (black) depict prediction metrics for a particular random state, number of downsampled reads, and *scAge* profiling parameter. Boxplots depict the median and the 1^st^ and 3^rd^ quartiles, with whiskers extending to 1.5x the interquartile range. Boxplots are colored based on the *scAge* parameter used for these predictions (i.e., the number of CpGs to profile in the likelihood computation). Upper plots in each panel depict the T statistic from Welch’s t-test used to quantify statistical significance between epigenetic age in fibroblasts and iPSC samples, with negative values indicating lower average epigenetic age in iPSC samples. Lower panels depict the one-tailed p-values associated with these t-tests. Red asterisk (*) highlights that significant p-values below 0.05 were reached. The dashed lines in the upper panels highlight T = 0, and the dashed lines in the lower panels depict nominal significance (p = 0.05).

Together, these data suggest that epigenetic age reversal by Yamanaka factor-based induction of pluripotency (conventional iPSC generation) can be assayed using low-pass approaches in combination with our clock framework, inviting further applications for the robust, inexpensive, and high-throughput evaluation of putative rejuvenation interventions.

### Low-coverage profiling uncovers the effect of genetic manipulations

In addition to dietary and reprogramming-based interventions, we were interested in testing if the integration of low-pass sequencing approaches with our *scAge* framework enables assessment of attenuated biological aging due to commonly studied genetic alterations. Some genetic manipulations, notably growth hormone receptor knockouts (GHRKO) and *Pit1* loss-of-function mutations (Snell dwarf), have previously shown lifespan-extending effects as well as decreased biological age when assessed by a blood epigenetic clock^10, 37, 38^.

We tested our approach on young GHRKO, Snell dwarf, and control mice (all independent from the Petkovich *et al.* reference set), and observed a trend towards age reduction, particularly when employing the Petkovich *et al.* models in samples with 100,000 reads and several thousand CpGs included in likelihood profiles (Fig. 7). The Thompson *et al.* models showed some nominal significance on GHRKO mice when 100,000 reads were downsampled, but this was not always consistent across random states. By selecting the best performing model based on benchmarking (100,000 reads per sample with the top 10,000 CpGs in each likelihood profile), we observed significant decreases in delta age for GHRKO mice compared to control mice across both reference sets, while the attenuated aging phenotype in Snell dwarf mice could only be significantly picked up by *scAge* running on the Petkovich *et al.*^10^ reference models (Extended Data Fig. 7, 8). This is not entirely unsurprising, as previous deep sequencing elastic-net clocks have also failed to pick up statistically significant differences in Snell dwarf models^11^. Interestingly, delta ages obtained with Thompson *et al.*-based *scAge* predictions were not centered around 0 (as is expected and was the case with Petkovich *et al*. models), reinforcing the likely presence of batch effects.

**Figure 7:**
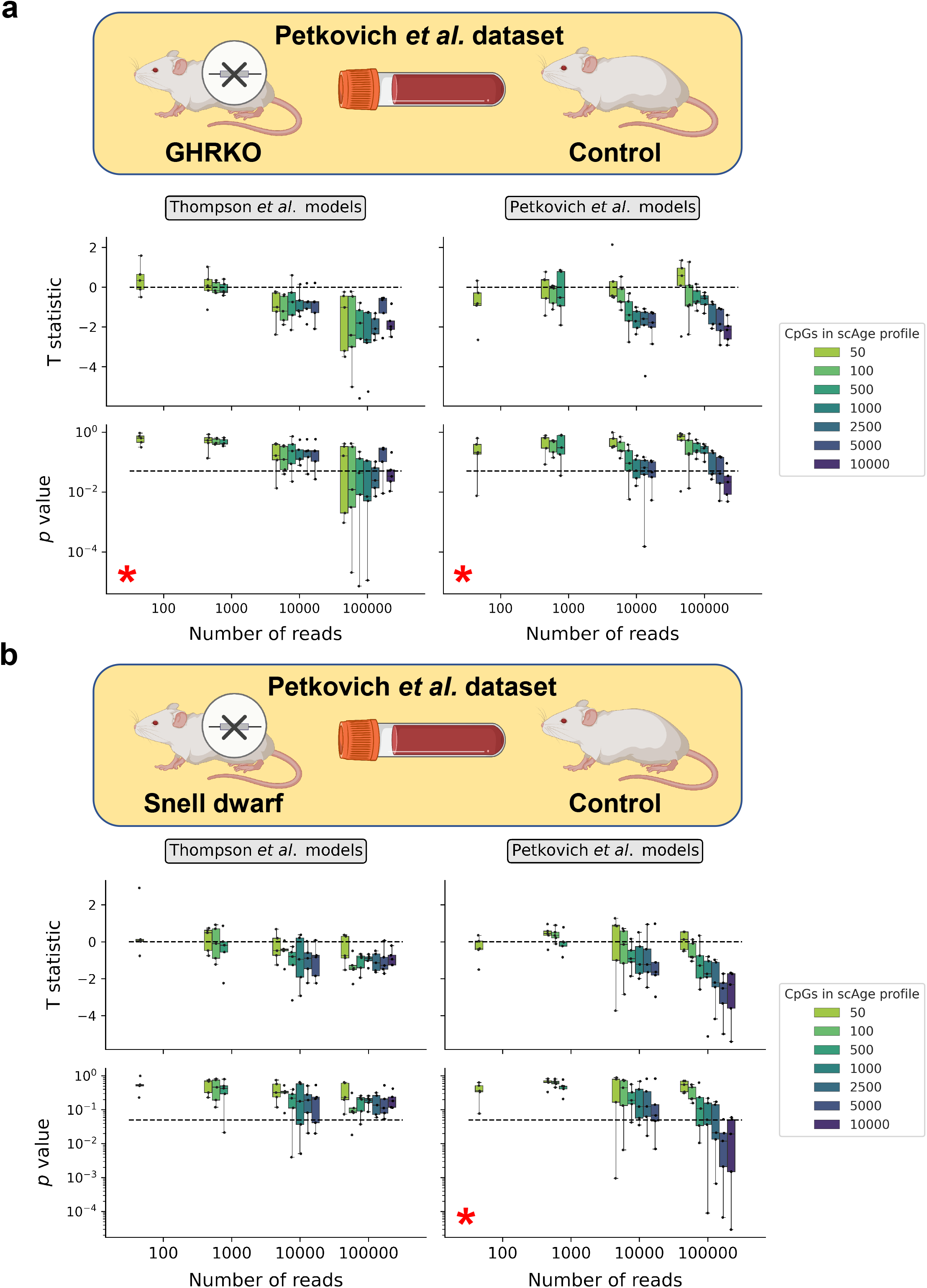
Low-coverage profiling tracks reduction in age from genetic manipulations. a-b) Experimental schematics (top), and boxplots (bottom) of statistical testing metrics based on epigenetic age measurements in (a) GRHKO (n = 15) and wildtype C57BL/6J x BALB/cByJ)/F2 (n = 11) blood samples of both sexes, aged 6 months and (b) Snell dwarf (n = 8) and wildtype (DW/J x C3H/HEJ)/F2 (n = 10) blood samples of both sexes, aged 6 months, from the Petkovich *et al.*^10^ study. Significance testing metrics using the Thompson *et al*. reference models are shown in the left panels, and those using the Petkovich *et al.* models are shown on the right. Individual dots (black) depict prediction metrics for a particular random state, number of downsampled reads, and *scAge* profiling parameter. Individual boxplots depict the median and the 1^st^ and 3^rd^ quartiles, with whiskers extending to 1.5x the interquartile range. Boxplots are colored based on the *scAge* parameter used for these predictions (i.e., the number of CpGs to profile in the likelihood computation). Upper plots in each panel depict the T statistic from Welch’s t-test used to quantify statistical significance between epigenetic age in genetically altered and wild-type samples, with negative values indicating lower epigenetic age in GHRKO/Snell dwarf samples. Lower panels depict the one-tailed p-values associated with these t-tests. Red asterisk (*) highlights that significant p-values below 0.05 were reached. The dashed lines in the upper panels mark T = 0, and the dashed lines in the lower panels highlight nominal significance (p = 0.05).

Altogether, we show that our new approach may discern the attenuated epigenetic aging effect brought on by genetic manipulations. Hence, low-pass sequencing combined with our computational age profiling platform opens avenues for large-scale genetic screening applications.

## Discussion

We report here the development and application of the epigenetic clock framework *scAge* to shallow bulk RRBS sequencing data, revealing robust performance at age prediction across independent murine blood datasets (Fig. 1-3). We randomly downsampled existing deep RRBS data, followed by epigenetic age profiling leveraging the *scAge* approach. As expected, predictions were stronger when the linear model reference set used came from the same study as the downsampled data, but independent reference datasets still performed well, suggesting that our approach should be generalizable to new collections of data.

Interestingly, we show that *scAge* combined with low-pass sequencing is amenable to reproducible profiling of biological age (Fig. 4). Our approach also enables tracking the attenuated biological aging effect brought on by calorie restriction (Fig. 5), as well as age reversal by induction of pluripotency through the Yamanaka factors in fibroblast lines from two tissues (Fig. 6). However, we note that we observe the most significant results for these interventions in the case where predictions are based on training data sourced from the same dataset as the downsampled data. This suggests the importance of acknowledging and accounting for batch effects in these analyses (Extended Data Fig. 3). Generation of additional bulk RRBS datasets amenable for training may shed more light on this phenomenon, and potentially enable universal application of our framework to other datasets. Additionally, we show that some genetic interventions—namely GHRKO and Snell dwarf mouse models—reduce biological age as predicted by *scAge*, but only under a more restrictive set of parameters and higher read counts (Fig. 7). Since the effect of caloric restriction can be picked up with high significance, we hypothesize that these slightly weaker results in genetic models may arise from application of a framework trained on C57BL/6J mice to different mouse strains; indeed, Snell dwarf mice and controls were of a (DW/J x C3H/HEJ)/F2 background, and GRHKO mice and controls were of a (C57BL/6J x BALB/cByJ)/F2 background.

A major advantage of our approach is the cost of sequencing. Compared to a typical RRBS protocol that yields >10^7^ reads per sample, our method at present requires only 10^4^-10^5^ reads, representing a 100-1,000-fold reduction in sequencing costs with appropriate multiplexing. An additional advantage of our method over those currently available is the direct use of isolated DNA samples for bisulfite sequencing, without the need to specifically enrich CpG sites of interest. This makes our method easily applicable to diverse data and sample sets, and does not require special expertise or reagents that are otherwise needed with methods relying on particular sets of pre-defined CpG sites.

While the framework we present here shows promise at accurate profiling of epigenetic age from low-pass sequencing data, there are a number of potential limitations that need to be acknowledged. First, while the approach employed in our study enables benchmarking and comparative analyses, it is not identical to high-throughput low-pass data, when more samples are multiplexed and sequenced in real-time. Additionally, our current method has so far only been applied in mice, although we suppose that identical or similar methods should be functional for low-cost epigenetic age profiling in human samples, as well as in other mammalian species. Our *scAge* approach also currently generalizes all methylation-age relationships as linear, which is likely not the most optimal mathematical reflection for the diverse set of age-related behaviors of individual CpGs. Lastly, while the use of different cytosines for age predictions in each sample enables *de novo* assessment of epigenetic aging (which could not otherwise be performed with conventional approaches), this reliance on diverse sets of CpGs may concurrently reduce the precision of predictions.

Overall, we leveraged our *scAge* framework for epigenetic age profiling in low-coverage bulk RRBS data. We assessed shallow data across a variety of read sets ranging from 10^2^ to 10^5^, revealing robust performance of our platform specifically when 10,000-100,000 reads are available per sample. We show that parameters of *scAge* have some impact on prediction metrics, although comparatively less than the number of reads sampled. Excitingly, we find that the *scAge* approach combined with shallow sequencing data enables prediction of biological age from as a little as 10,000 reads in standard blood samples of C57BL/6J mice across two independent datasets. Additionally, we report that our method may be amenable to identify the effects of some longevity and rejuvenation interventions—such as calorie restriction or iPSC reprogramming—on biological aging trajectories. This suggests our approach may be particularly useful to validate and identify new longevity interventions.

While epigenetic age profiling is becoming more mainstream, current methods remain prohibitively expensive for routine epigenetic testing or for large-scale assessment of biological age in human cohorts or in model systems. Compared to deep RRBS sequencing and conventional epigenetic clock analyses, this application of *scAge* reflects an up to 1000-fold reduction in sequence read numbers needed to assess epigenetic age. As such, this approach, in tandem with parallel library preparation workflows and optimized multiplexing strategies, may be applicable to large-scale data such as genetic screens, interventions studies and biobanks. Accordingly, these approaches may have important implications for clinical testing and in consumer settings. Overall, we present a novel computational approach which may enable accurate, rapid and cost-effective epigenetic age profiling from low-pass sequencing approaches.

## Methods

### Animal resources and mouse models

Mice originating in the Petkovich *et al*. study^10^, with the exception of Snell dwarf, GHRKO models and 3-5-week-old C57BL/6 mice, were obtained from the NIA Aged Rodent Colony. The oldest mice obtained from NIA were 32 months; to obtain mice 34 and 35 months of age, 32-month-old mice were aged at Brigham & Women’s Hospital (BWH) for 2 and 3 months, respectively. With the exception of mice at the 34-and 35-month time points, blood from all mice used was collected within 2 weeks of arrival from NIA. Snell dwarf and GHRKO models and their controls were housed at the University of Michigan. The 3-5-week-old C57BL/6 mice were bred at BWH, Harvard Medical School, from parents obtained from NIA. Calorie restriction started at 14 weeks of age and continued until the time the animals were sacrificed. For all other animals in the Petkovich *et al*. study, food was provided *ad libitum*. All mouse data used from the Thompson *et al*. study^27^ were from *ad libitum* C57BL/6J mice housed at UCLA, which ranged in age from 2-21 months.

### Downsampling methylation data

Methylation data were obtained from the GEO database for both the Petkovich *et al.*^10^ and Thompson *et al.*^27^ datasets, with accession numbers GSE80672^10^ and GSE120132^27^, respectively. Metadata was also downloaded from the GEO database using the *GEOparse* 2.0.3 package. Methylation sequences were mapped to the mm10/GRCm38 mouse genome, and methylation information was extracted by the original authors with Bismark^39^ in the case of the Petkovich *et al.* data, and BS-Seeker 2^40^ in the case of the Thompson *et al.* data. We further filtered processed methylation data to include only CpGs on autosomic chromosomes, in order to partially mitigate the effect of sex on predictions. Individual CpG reads (whether methylated or unmethylated) were concatenated into lists in a randomized order to prevent location bias from affecting downstream predictions. We then selected a defined number of reads ranging from 100 to 100,000, each covering mostly unique CpGs, with some overlap when a larger number of reads were sampled. Methylation at CpGs in downsampled data was calculated as the mean of all methylation reads for that CpG in a particular subsample. Given the low number of reads, this most often produced binary methylation values (0, unmethylated; 1, methylated), with some additional fractional values in between this range that arose when multiple reads covered a single CpG. In order to produce random subsamples, we used the *sample* function from the *random* Python library with five different reproducible seeds, enabling the generation of distinct assortments of CpGs for the same sample.

### Generating scAge reference models

To create linear regression models enabling epigenetic age profiling via the *scAge* framework, we utilized deeply sequenced training data from the Petkovich *et al.*^10^ and Thompson *et al.*^27^ studies. Specifically, we filtered for only standard *ad libitum* blood C57BL/6J samples, resulting in dataframes with n *=* 153 and n *=* 50 samples for the Petkovich *et al.*^10^ and Thompson *et al*.^27^ studies, respectively. For each individual sample, only CpG sites covered 5 times or greater were retained, while other CpG sites were marked as missing. This filtering ensured high resolution estimation of methylation proportions in a particular sample, which is an integral component of the *scAge* training framework. Next, we progressively intersected all samples using an outer join methodology, capturing all CpGs covered 5x or more in at least one sample in the dataset. In order to select only CpGs which were consistently deeply covered across samples (facilitating accurate linear regression modeling), we removed CpG sites for which more than 10% of samples had missing values after depth filtration.

By applying these methods to both maximize CpG coverage and methylation value accuracy, we ultimately arrived at a table of 1,918,766 CpGs across 153 samples in the Petkovich *et al.*^10^ dataset and 1,202,751 CpGs across 50 samples in the Thompson *et al.*^27^ dataset. Of note, the Petkovich *et al.*^10^ dataset contained both negative and positive strand CpGs, while the Thompson *et al.*^27^ dataset contained only positive strand CpGs. This is a result of the processing methods used, wherein Thompson *et al*.^27^ concatenated positive and negative strand reads to the positive strand to increase confidence in the methylation values while decreasing the total feature space. This difference in processing, combined with significant batch effects, may help to explain the deviations that we observe when comparing predictions based on each reference model.

### Applying the scAge prediction framework

To assess epigenetic age in low-coverage bulk data, we could not utilize conventional mouse methylation clocks, nor could we train novel methylation clocks based on classical elastic-net regression approaches. This is entirely due to the primary limitations of downsampled data: the lack of consistent CpG coverage across samples, coupled with a primarily binary data type. In order to overcome these fundamental constraints, we made use of the recently developed *scAge* framework, which enables accurate epigenetic age profiling in single cells^26^. Single-cell data features notoriously sparse, binary methylation profiles as a function of current limitations in sequencing protocols, which happen to heavily resemble downsampled bulk RRBS data. However, while the binary nature of single-cell methylation profiles is consistent with the biology underlying this data, bulk methylation profiles often contain fractional methylation values within the unit region, representing the proportion of cells in a specific sample that are methylated at a particular cytosine. Of note, these changes in fractional DNA methylation across bulk samples have been particularly key to methylation clock development in the past.

Practically, downsampled data were first intersected on an individual basis with a particular reference set. Next, of the common CpG sites between any subsample and the reference, a defined number (varied as a parameter in the analyses) of the most age-associated CpG sites—based on the magnitude of Pearson correlations—was chosen. This enabled testing distinct combinations of three critical parameters at once: the number of downsampled reads, the number of CpGs included in the *scAge* likelihood profile, and the particular random subsample. Among selected age-associated CpG sites, the distance between the observed methylation value and the linear regression estimate for a particular cytosine was treated as a probability measure. In essence, we reasoned that the closer the observed methylation value was to the linear regression estimate for a particular age, the higher the probability of observing this methylation state at this age. Hence, we subtracted the distance between the observed value and the linear estimate from 1, then proceeded to take the natural logarithm of the resulting difference to circumvent underflow errors in downstream processing. Once all relevant CpG sites were assayed for a particular age, we summed logarithmic probabilities together for the entire profile (equivalent to taking the product of all probabilities), generating a single value describing the likelihood of observing a particular limited methylome for each age. An age-likelihood distribution was hence created, mapping probabilities between −20 months and 60 months in high-resolution intervals of 0.1 months, ultimately enabling prediction of epigenetic age based on maximum likelihood estimation.

### Computational and statistical analyses

All analyses were performed using Python 3.9.2, running with *numpy* 1.20.2 and *pandas* 1.2.4 for mathematical computing and data wrangling. Figures were generated using *matplotlib* 3.4.1 in combination with *seaborn* 0.11.1. Welch’s one-tailed t-test assuming unequal variances, implemented in *statannot* 0.2.3 and *scipy* 1.6.3, was used to perform statistical tests between groups. Pearson correlation coefficients and associated two-tailed p-values were computed using the *pearsonr* function implemented in the *scipy* package. Linear regressions underlying the *scAge* framework were computed with the *LinearRegression* function implemented in *scikit-learn* 0.24.2. Delta ages were computed as the difference between epigenetic age and chronological age.

## Acknowledgments

We are grateful to Csaba Kerepesi, Patrick Griffin, Yan Hu, Marco Mariotti, Anastasia Shindyapina, Sun Hee Yim, Bohan Zhang, Benjamin Barre, Kejun “Albert” Ying, Jesse Poganik, Sang-Goo Lee, and Didac Santesmasses for helpful discussion. This work was supported by NIA grants to VNG. Some figures were created with BioRender.com.

## Author Contributions

AT performed all analyses in the manuscript and developed the modified *scAge* framework. AT and VNG conceived the study and wrote the manuscript. VNG supervised the work.

## Data Availability

All data used in this work were obtained from publicly available repositories. Bulk methylation data used for reference model training and downsampling was downloaded from GEO under accession numbers GSE120132^27^ and GSE80672^10^.

## Code Availability

Source code for the modified *scAge* framework will be made available on GitHub.

---

**Extended Data Figure 1:**
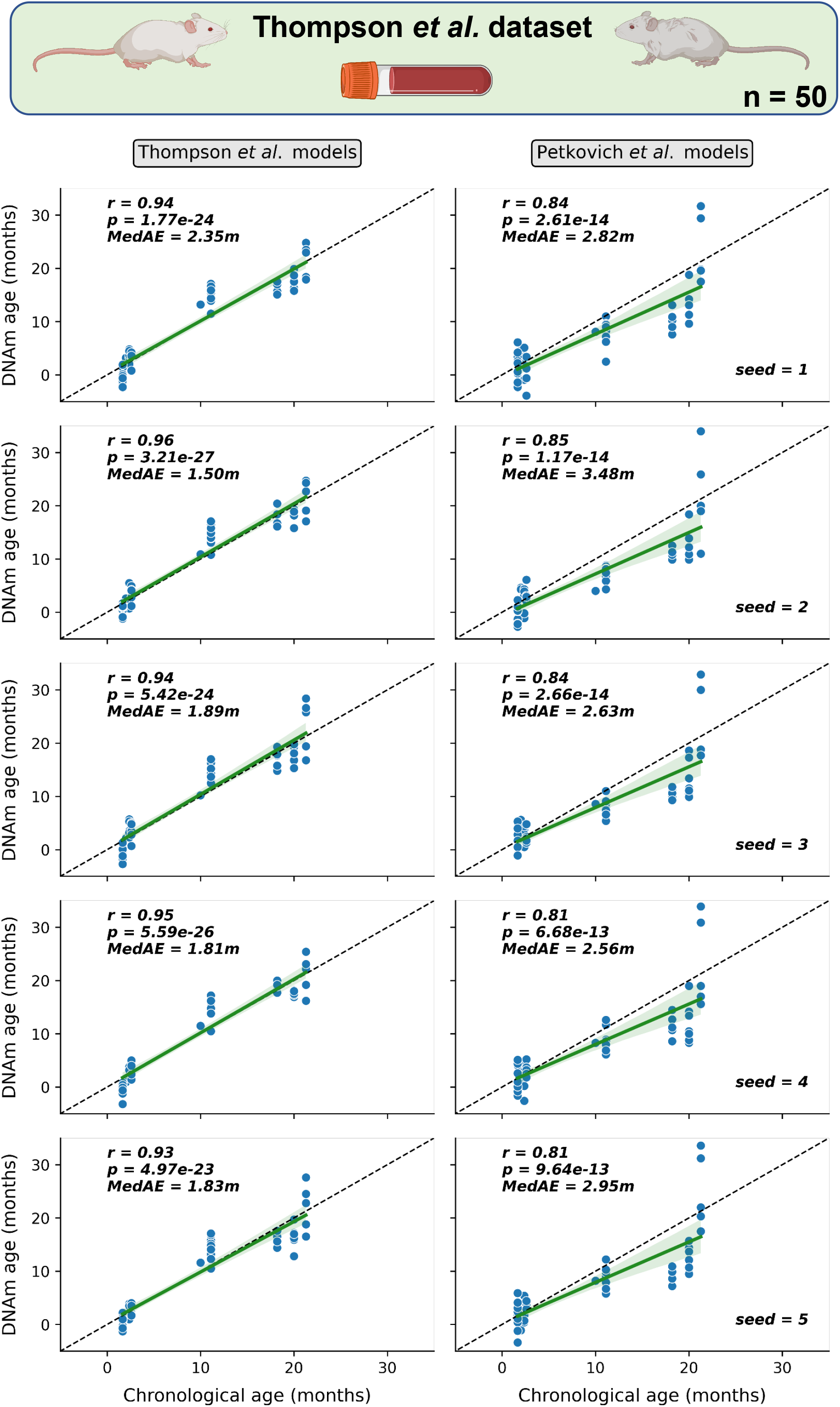
Low-pass epigenetic age predictions in the Thompson *et al.* blood dataset. Scatterplots of epigenetic age predictions for C57BL/6J blood samples in the Thompson *et al.* dataset (n *=* 50). Panels on the left depict predictions using Thompson *et al.* reference models, while panels on the right show predictions using regression models computed with Petkovich *et al.* data. The particular random downsampling seed used is shown in the bottom right and corresponds across training datasets. The Pearson correlation coefficient (*r*), the associated two-tailed p-value (*p*), and the median absolute error (*MedAE*) is shown for each plot. Data shown are from the best performing set of parameters (100,000 reads and 1,000 CpGs included in the likelihood profile), based on benchmarking (see Figure 2). Regression lines (green) are shown with 95% confidence intervals (light green). The dashed lines represent the identity between epigenetic (DNAm) and chronological age.

**Extended Data Figure 2:**
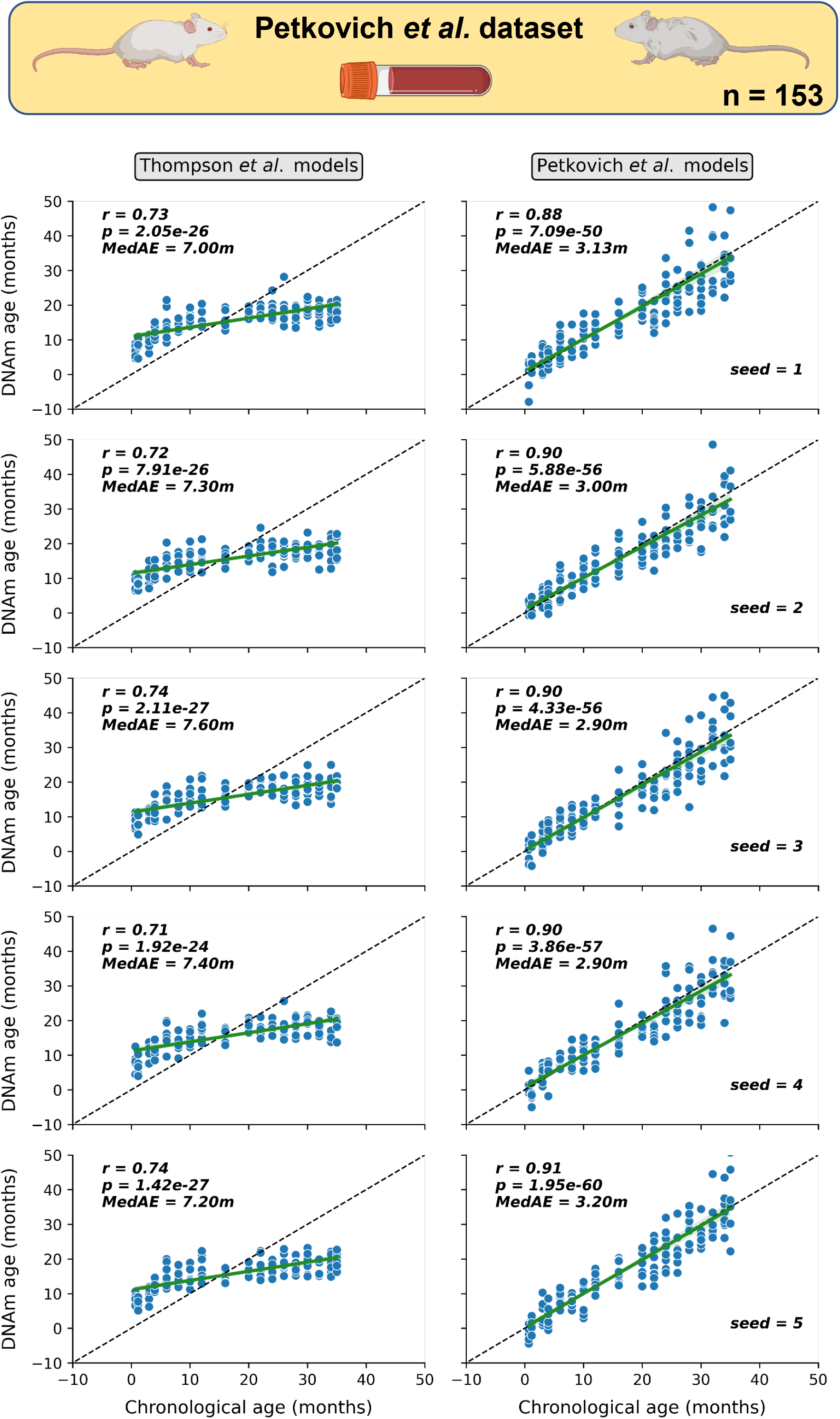
Low-pass epigenetic age predictions in the Petkovich *et al.* blood dataset. Scatterplots of epigenetic age predictions for C57BL/6J blood samples in the Petkovich *et al.* dataset (n *=* 153). Panels on the left depict predictions using Thompson *et al.* reference models, while panels on the right show predictions using Petkovich *et al.* regression models. The particular random downsampling seed used is shown in the bottom right and corresponds across training datasets. The Pearson correlation coefficient (*r*), the associated two-tailed p-value (*p*), and the median absolute error (*MedAE*) is shown for each plot. Data shown are from the best performing set of parameters (100,000 reads and 500 top CpGs included in the likelihood profile), based on benchmarking (see Figure 3). Regression lines (green) are shown with 95% confidence intervals (light green). The dashed lines represent the identity between epigenetic (DNAm) and chronological age.

**Extended Data Figure 3:**
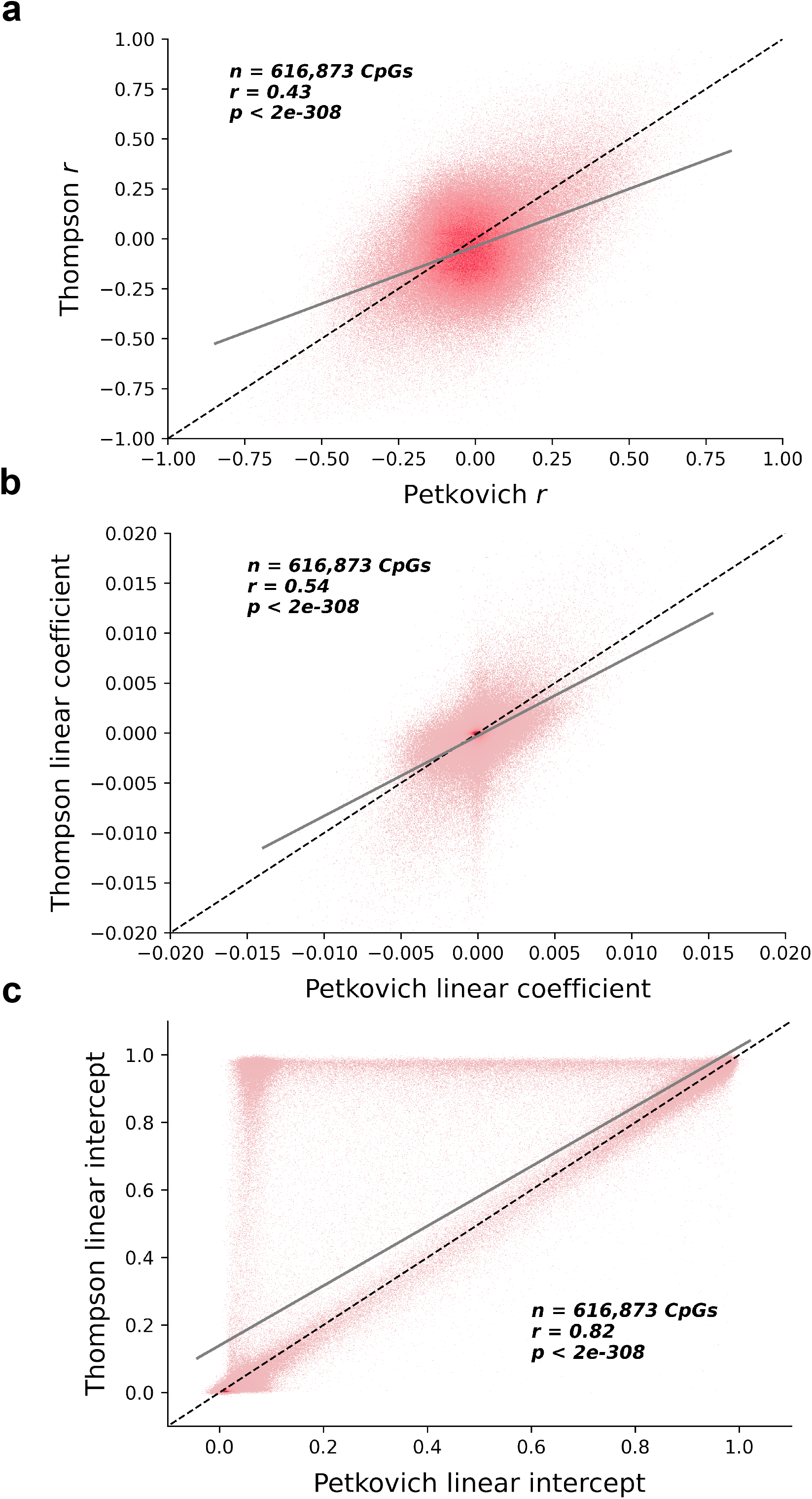
Comparison of linear association metrics with age in the Thompson *et al.* and Petkovich *et al.* datasets. Density plots depicting the correspondence of (a) Pearson correlation coefficients, (b) linear regression coefficients, and (c) linear regression intercepts between training data from Petkovich *et al.* (x-axis) and Thompson *et al.* (y-axis). The number of common CpGs between both datasets is shown (*n*), as well as the Pearson correlation coefficient (*r)* and the associated two-tailed p-value. Dashed lines represent identity between a metric in the Petkovich *et al.* dataset and that same metric in the Thompson *et al*. dataset. Regression lines (grey) are shown with 95% confidence intervals (light grey).

**Extended Data Figure 4:**
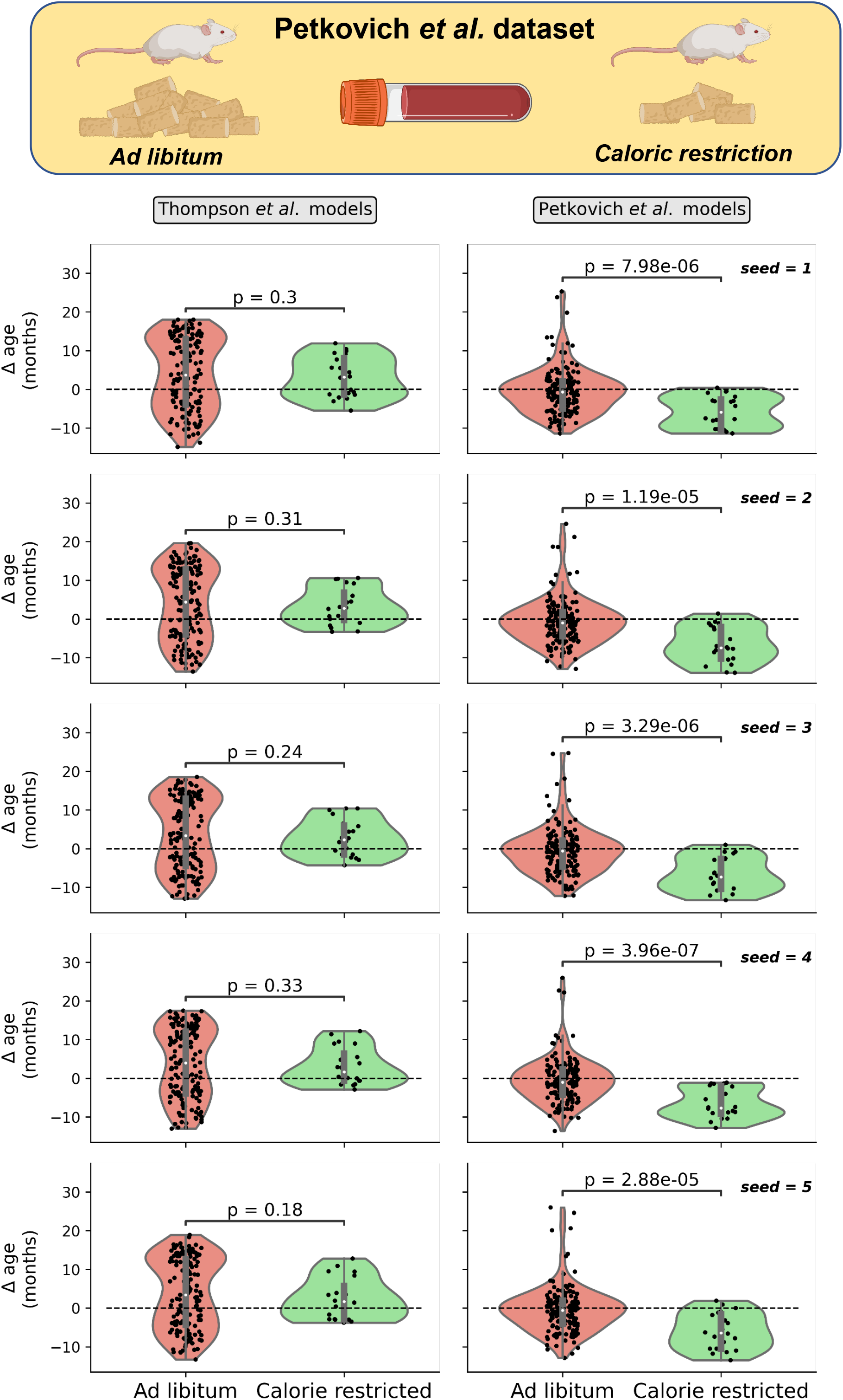
Decreased delta age in calorically-restricted samples. Violin plots of delta age (epigenetic age – chronological age) for *ad libitum* (n *=* 153, red) and calorically-restricted (n *=* 20, green) male C57BL/6J blood samples in the Petkovich *et al.* dataset. Panels on the left depict predictions using Thompson *et al.* reference models, while panels on the right show predictions leveraging Petkovich *et al.* data. The particular random downsampling seed used is shown in the top right. The p-values depicted are derived from Welch’s one-tailed t-test (assuming unequal variances). Data shown are from the best performing set of parameters (100,000 reads per sample and 2,500 CpGs in the likelihood profile) based on benchmarking. Inner boxplots depict the median and the 1^st^ and 3^rd^ quartiles, with whiskers extending to 1.5x the interquartile range. Violin plots show the kernel density estimation of the data. Dashed lines highlight a delta age of 0.

**Extended Data Figure 5:**
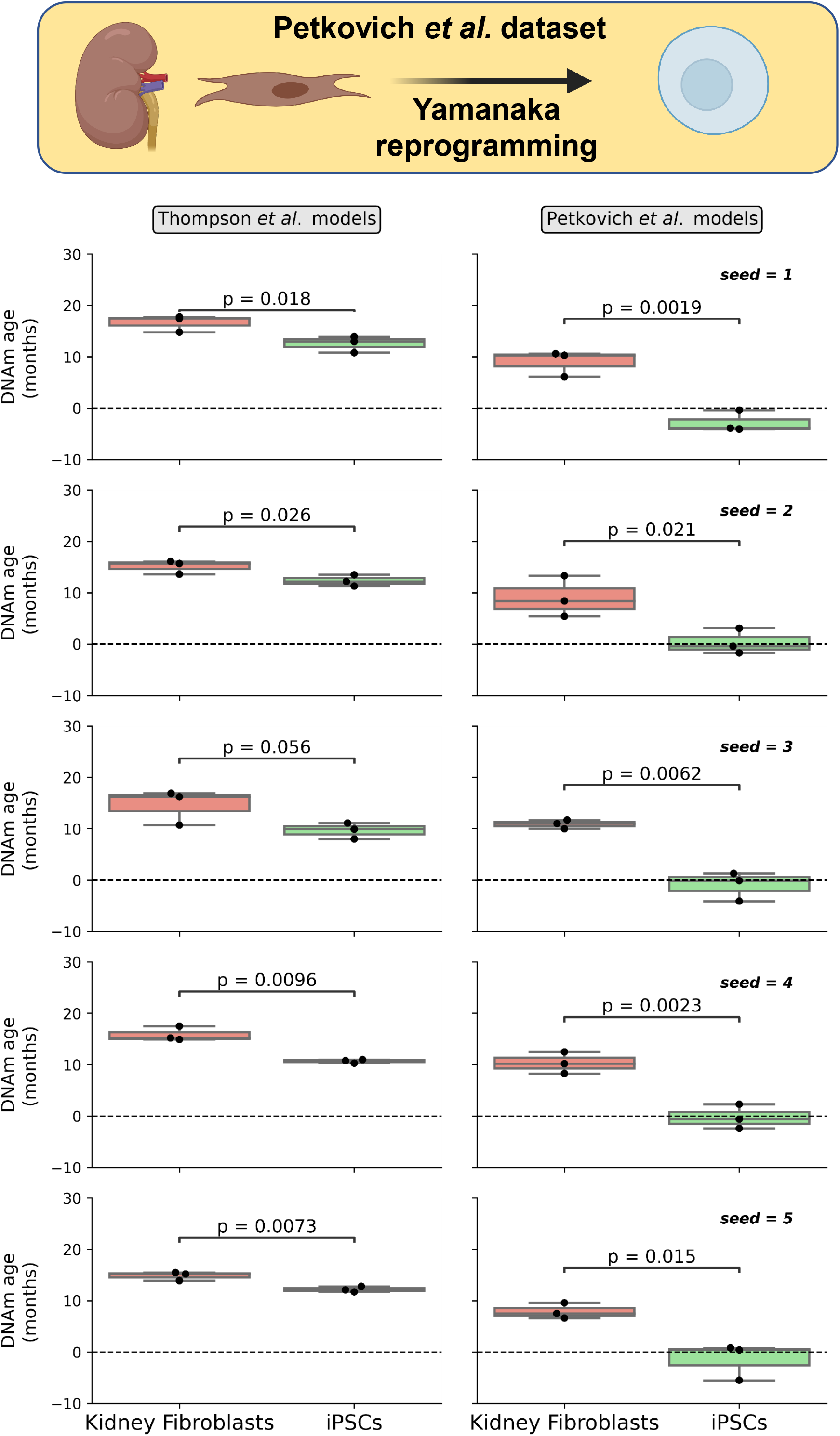
Age reversal assessed by low-pass *scAge* in renal fibroblasts and derived iPSCs. Box plots of epigenetic age for kidney fibroblasts (n *=* 3, red) and kidney-derived induced pluripotent stem cells (iPSCs, n = 3, green) from C57BL/6J mice in the Petkovich *et al.*^10^ dataset. Panels on the left depict predictions using Thompson *et al.* reference models, while panels on the right show predictions using Petkovich *et al.* regression models. The particular random downsampling seed used is shown in the top right and corresponds across training datasets. The p-values depicted are derived from Welch’s one-tailed t-test (assuming unequal variances). Data shown are from the best performing set of parameters (100,000 reads per sample and the top 500 age-associated CpGs in the likelihood profile). Boxplots depict the median and the 1^st^ and 3^rd^ quartiles, with whiskers extending to 1.5x the interquartile range. Dashed lines indicate an epigenetic (DNAm) age of 0.

**Extended Data Figure 6:**
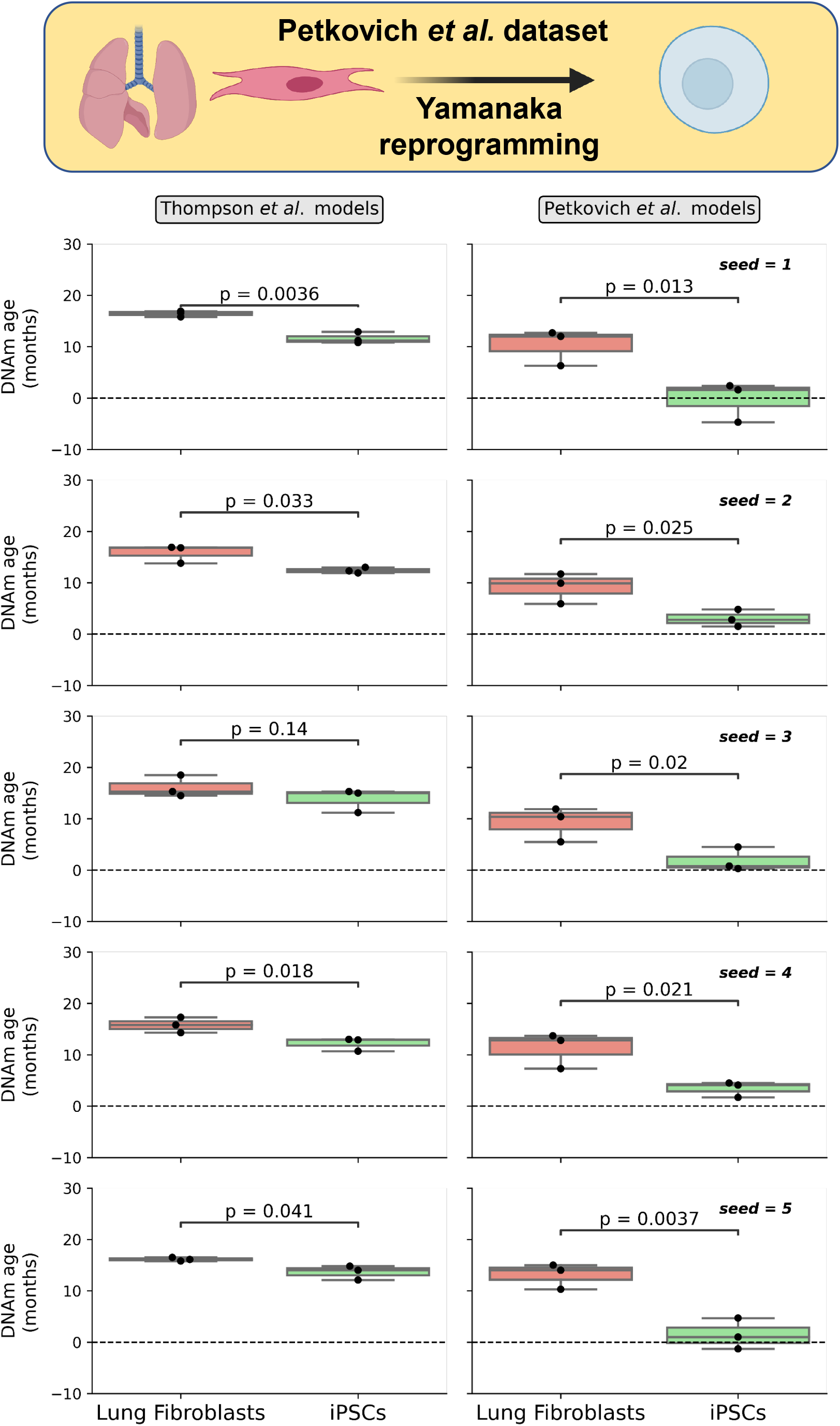
Age reversal assessed by low-pass *scAge* in lung fibroblasts and derived iPSCs. Box plots of epigenetic age for lung fibroblasts (n *=* 3, red) and lung-derived induced pluripotent stem cells (iPSCs, n = 3, green) from C57BL/6J mice in the Petkovich *et al.*^10^ dataset. Panels on the left depict predictions using the Thompson *et al.* training dataset, while panels on the right show predictions using the Petkovich *et al.* regression models. The particular random seed used is shown in the top right and corresponds across training datasets. The p-values depicted are derived from Welch’s one-tailed t-test (assuming unequal variances). Data shown are from the best performing set of parameters (100,000 reads per sample and the top 500 age-associated CpGs in the likelihood profile). Boxplots depict the median and the 1^st^ and 3^rd^ quartiles, with whiskers extending to 1.5x the interquartile range. Dashed lines indicate an epigenetic (DNAm) age of 0.

**Extended Data Figure 7:**
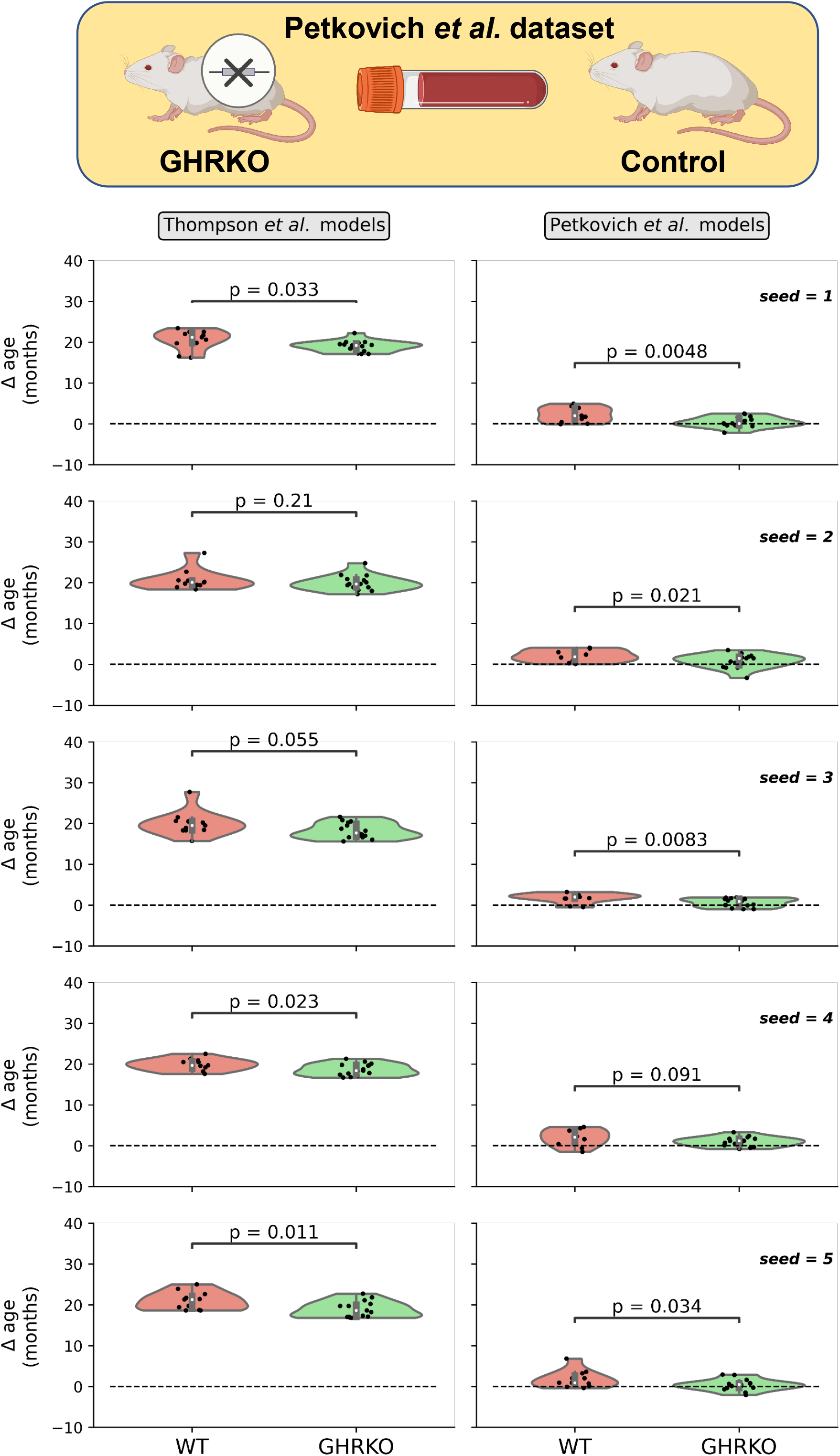
Decreased delta age in GHRKO samples. Violin plots of delta age (epigenetic age – chronological age) for wild type (n *=* 11, red) and growth hormone receptor knockout (GHRKO, n = 15, green) samples from (C57BL/6J x BALB/cByJ)/F2 mice in the Petkovich *et al.*^10^ dataset. Panels on the left depict predictions using Thompson *et al.* reference models, while panels on the right show predictions using Petkovich *et al.* regression models. The particular random downsampling seed used is shown in the top right and corresponds across training datasets. The p-value depicted is derived from Welch’s one-tailed t-test (assuming unequal variances). Data shown are from the best performing set of parameters (100,000 reads per sample with the top 10,000 CpGs included in the *scAge* profile). Inner boxplots depict the median and the 1^st^ and 3^rd^ quartiles, with whiskers extending to 1.5x the interquartile range. Violin plots show the kernel density estimation of the data. Dashed lines indicate a delta age of 0.

**Extended Data Figure 8:**
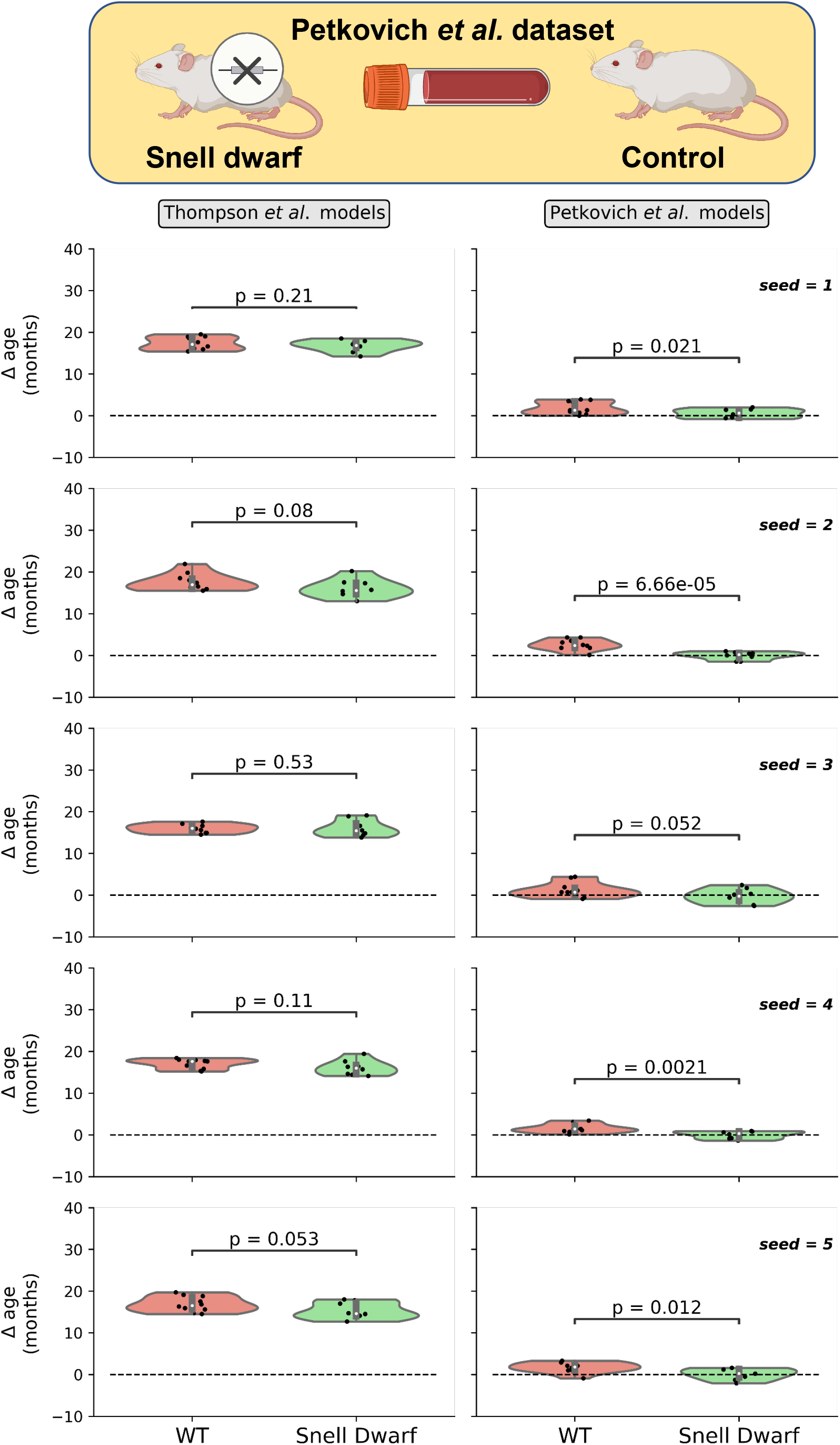
Decreased delta age in Snell dwarf samples. Violin plots of delta age (epigenetic age – chronological age) for wild type (n *=* 10, red) and Snell Dwarf (n = 8, green) samples from (DW/J x C3H/HEJ)/F2 mice in the Petkovich *et al.*^10^ dataset. Panels on the left depict predictions using Thompson *et al.* reference models, while panels on the right show predictions using Petkovich *et al.* models. The particular random seed used is shown in the top right and corresponds across training datasets. The p-value depicted is derived from Welch’s one-tailed t-test (assuming unequal variances). Data shown are from the best performing set of parameters (100,000 reads per sample with the top 5,000 CpGs included in the profile). Inner boxplots depict the median and the 1^st^ and 3^rd^ quartiles, with whiskers extending to 1.5x the interquartile range. Violin plots show the kernel density estimation of the data. Dashed lines indicate a delta age of 0.

